# SOX2 Regulation by Hedgehog Signaling Controls Adult Lingual Epithelium Homeostasis

**DOI:** 10.1101/269522

**Authors:** David Castillo-Azofeifa, Kerstin Seidel, Lauren Gross, Belkis Jacquez, Ophir D. Klein, Linda A. Barlow

**Author notes:** current address: Program in Craniofacial Biology and Department of Orofacial Sciences, University of California San Francisco, San Francisco, CA 94131, USA. current address: Department of Discovery Oncology, Genentech, Inc., South San Francisco, CA.

## Abstract

The adult tongue epithelium is continuously renewed from epithelial progenitor cells, and this process relies on intact Hedgehog (HH) signaling. In mice, inhibition of the HH pathway using Smoothened antagonists (HH pathway inhibitors or HPIs) leads to taste bud loss over a span of several weeks. Previously, we demonstrated that overexpression of Sonic Hedgehog (SHH) in lingual epithelial progenitors induces formation of ectopic taste buds accompanied by locally increased SOX2 expression, consistent with the hypothesis that taste bud differentiation depends on SOX2 downstream of HH. To test this idea, we inhibited HH signaling by treating SOX2-GFP mice with HPI and found a rapid and drastic decline in SOX2-GFP expression in taste progenitors and taste buds. Using a conditional Cre-lox system to delete *Sox2*, we found that loss of SOX2 blocks differentiation of both taste buds and non-taste epithelium that comprises the majority of the tongue surface; progenitor cells increase in number at the expense of differentiated taste cells and lingual keratinocytes. In contrast to the normal pattern of basally restricted proliferation, dividing cells are overabundant, disorganized and present in suprabasal epithelial layers in *Sox2* deleted tongues. Additionally, SOX2 loss in taste progenitors leads non-cell autonomously to rapid loss of taste bud cells via apoptosis, dramatically shortening taste cell lifespans. Finally, when *Sox2* is conditionally deleted in mice with constitutive overexpression of SHH, ectopic taste buds fail to form and endogenous taste buds disappear; instead, robust hyperproliferation takes over the entire lingual epithelium. In sum, our experiments suggest that SOX2 functions downstream of HH signaling to regulate lingual epithelium homeostasis.

## Introduction

In mammals, the adult lingual epithelium can be categorized into non-taste and taste components. The majority of the tongue surface is covered by keratinized, non-taste epithelium made up of mechanosensory filiform papillae, which are curved, spinous-shaped structures with small mesenchymal cores (Hume and Potten, 1976). The more complex taste epithelium consists of collections of neuroepithelial taste cells organized within taste buds, which in turn lie in specialized taste papillae on the tongue surface. In rodents, fungiform papillae (FFP) are arrayed on the anterior two-thirds of the tongue, interspersed among filiform papillae of the non-taste epithelium. Each rodent FFP houses a single apical taste bud surrounded by keratinocytes that make up the papilla walls that in turn surround a mesenchymal core. Murine taste buds comprise ~60-100 fusiform taste receptor cells responsible for detecting, transducing and transmitting to the brain the five basic tastes, i.e. salty, sweet, bitter, umami/savory and sour (Chaudhari and Roper, 2010).

Both non-taste and taste epithelium are continually renewed from basally located progenitor cells. Previous tritiated thymidine studies in mice show that the non-taste epithelium has a turnover rate of 5-7 days; this is one of the fastest renewing tissues in mammals, only slightly slower than the pace of renewal of the epithelial lining of the intestine (3-5 days) (Hume and Potten, 1976; Liu et al., 2012; Barker, 2014). By contrast, similar approaches to define taste cell life span concluded these cells are longer lived, with a median lifespan of 10-14 days (Beidler and Smallman, 1965; Farbman, 1980; Delay et al., 1986), although some taste bud cells persist up to 44 days (Hamamichi et al., 2006; Perea-Martinez et al., 2013).

More recent studies using inducible genetic systems to lineage label cytokeratin 5 and 14 (K5 and K14) basal epithelial cells in the tongue have demonstrated that this population comprises bipotential progenitors (Okubo et al., 2009; Gaillard et al., 2015). K5/14^+^ cells located basally in the non-taste epithelium are progenitor cells that give rise to differentiated Keratin (K) 13^+^ keratinocytes. K13^+^ cells contribute to subrabasal epithelial layers of filiform and FF papillae, and are ultimately shed at the tongue surface (Iwasaki et al., 2006; Okubo et al., 2009); this lineage progression resembles that of the interfollicular epidermis of the skin (Winter et al., 1990). In taste epithelium, the K5/14^+^ cells situated immediately adjacent to each bud are referred to as perigemmal cells. These mitotically active progenitors generate cells that exit the cell cycle, enter taste buds, and become immediate taste cell precursors (also known as type IV cells) located basally in each bud. Ultimately these intragemmal cells, i.e., inside taste buds, differentiate into mature taste cells within 2.5-3 days of their last cell division (Cho et al., 1998; Hamamichi et al., 2006; Miura et al., 2006; Nguyen and Barlow, 2010; Perea-Martinez et al., 2013; Miura et al., 2014; Barlow, 2015; Barlow and Klein, 2015). Besides contributing to taste buds, K5/K14^+^ progenitor cells within FFP replenish themselves and provide keratinocytes to the region of the taste pore and adjacent, K13^+^ non-taste epithelium of the taste papillae (Okubo et al., 2009). The intrinsic differences between non-taste and taste epithelial cell lineages suggest differential molecular regulation of cell fates in these two tissue compartments.

One molecular regulator of cell renewal in many epithelia is the Sonic hedgehog (SHH) pathway. SHH is expressed by postmitotic taste precursor cells (type IV cells) (Miura et al., 2014), while mitotically active K5/K14^+^ progenitors surrounding each bud express the SHH target genes, *Ptch1* and *Gli1*, suggesting SHH signals from within the bud to adjacent progenitors, to regulate proliferation and/or taste cell differentiation (Miura et al., 2001; Miura, 2003; Miura and Barlow, 2010; Miura et al., 2014). HH signaling-dependent cancers, such as basal cell carcinomas, are frequently treated with HPIs to inhibit constitutive HH pathway activation (Rubin and de Sauvage, 2006; Ng and Curran, 2011; Wong and Dlugosz, 2014). Although these chemotherapeutics efficiently target tumors, patients experience distiburbingly altered taste sensation (LoRusso et al., 2011; Tang et al., 2012; Rodon et al., 2014). Moreover, in mice, HPIs lead to taste bud loss, as well as loss of taste nerve responses (Kumari et al., 2015; Yang et al., 2015; Kumari et al., 2017), indicating that HH signaling is required for taste cell renewal. In fact, we have recently shown that SHH functions to promote taste cell differentiation but is not required for perigemmal progenitor proliferation.

Another important regulator of taste bud differentiation is SOX2, which belongs to the family of SRY-related HMG box transcription factors that are critical for cell fate determination during development and stem cell maintenance/differentiation in many adult tissues (Arnold et al., 2011; Liu et al., 2012). The importance of SOX2 during embryonic taste development was first demonstrated by failure of *Sox2* hypomorphic mutants to generate FF taste buds (Okubo et al., 2006). In the adult tongue, SOX2 is expressed at low levels by K14^+^ cells, and *Sox2* genetic lineage tracing confirms that SOX2^+^ basal keratinocytes also function as bipotential stem cells for taste and non-taste lingual epithelium (Ohmoto et al., 2017). Further, SOX2 is robustly expressed in perigemmal K14^+^ taste bud progenitors, basal cells within taste buds and in a subset of mature taste receptor cells (Okubo et al., 2006; Suzuki, 2008; Ohmoto et al., 2017), suggesting that SOX2 may be key to proper taste cell differentiation from progenitors. Interestingly, overexpression of SHH in K14^+^ progenitors leads to formation of ectopic taste buds, which are associated with increased SOX2 expression in epithelial cells surrounding and within ectopic buds (Castillo et al., 2014). These results suggested the testable hypothesis that renewal of adult lingual epithelium is positively regulated by HH signaling, which in turn requires downstream SOX2 function.

Here, we test this idea by assessing the impact of HH pathway inhibition on SOX2 expression using HhAntag, a Hedgehog pathway inhibitor (HPI) that blocks the HH effector Smoothened (SMO) (Yauch et al., 2008). Additionally, we genetically delete *Sox2* (SOX2cKO)(Shaham et al., 2009) or pair SOX2cKO with SHH over-expression (SHH-YFPcKI)(Castillo et al., 2014) in K14+ progenitors to explicitly test if SOX2 is required for taste cell differentiation. We find that pharmacologic inhibition of the HH pathway, which blocks the differentiation program of taste buds (Castillo Azofeifa et al., 2017), also leads to downregulation of SOX2-GFP in taste bud progenitors and taste buds. Further we show that SOX2 function in lingual progenitors is required broadly for lingual epithelial cell maintenance; in SOX2cKO mice, K14+ progenitors fail to differentiate and proliferate inappropriately. Unexpectedly, we also find that SOX2 function in progenitors is required non-cell autonomously for survival of differentiated taste bud cells, as taste cells rapidly undergo apoptosis when *Sox2* is deleted only from progenitors. Finally, loss of SOX2 abrogates the ability of SHH to induce ectopic taste buds; instead, SHH overexpression in SOX2cKO epithelium results in hyperproliferation of basal epithelial cells, suggesting that, in the absence of SOX2, SHH switches from a pro-taste differentiation signal to a robust mitogen.

## Results

### Adult taste buds are mildly but significantly affected within 1 week of HhAntag treatment

Others and we have found that taste buds are significantly reduced after 21 days of HPI treatment (Kumari et al., 2015; Yang et al., 2015; Castillo-Azofeifa et al., 2017; Kumari et al., 2017); during 2-4 weeks of drug exposure, typically appearing FFP and their taste buds (see Fig. 1A) are gradually lost as the number of atypical, i.e., degenerating, FFP taste buds (see Fig. 1B, Oakley et al., 1990; Nagato et al., 1995) increases (Kumari et al., 2015, Castillo Azofeifa et al., 2017). However, because differentiation of taste progenitors into new taste cells takes ~3 days from their last division, we hypothesized that HPIs would affect taste bud renewal well in advance of taste bud loss. Thus, we looked for the shortest time of drug treatment that revealed an effect on taste bud maintenance. Using Keratin (K)8 immunostaining to mark mature taste buds in FFP (Fig 1A)(Knapp et al., 1995), we found that typical FFP taste bud number and size did not differ from controls after 3 days of drug (Fig 1C, D); the number of atypical FFP also was not increased after 3 days of HhAntag (Fig. 1C). By 7 days of HhAntag treatment, typical FFP number was minimally but not significantly decreased, a trend that was similar for taste bud size (Fig. 1E, F). However, atypical FFP number increased significantly in drug-treated mice compared to controls, suggesting that taste bud maintenance is affected within 7 days of HhAntag treatment.

**Figure 1.**
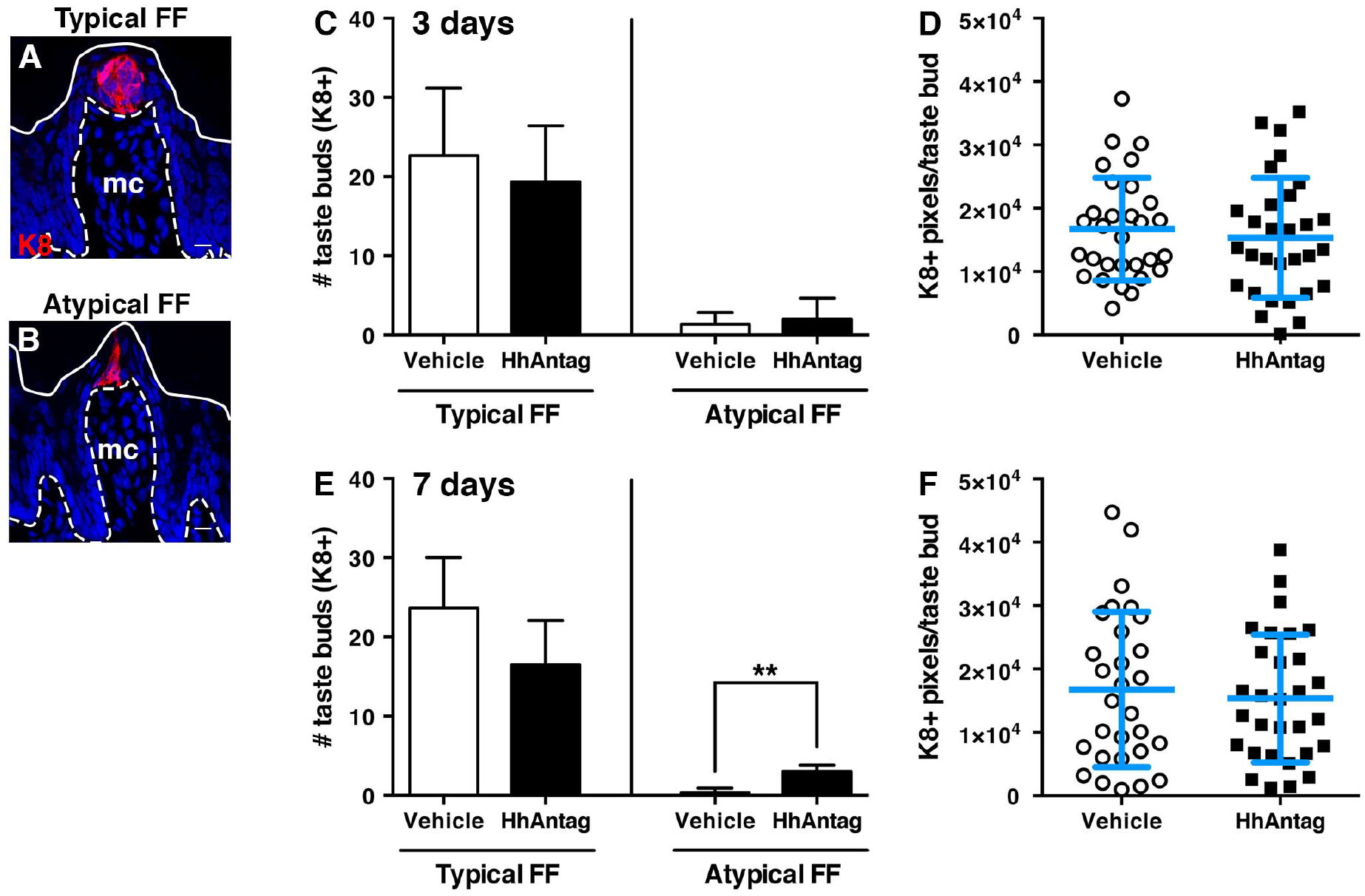
Adult taste buds are mildly but significantly affected after 1 week of HhAntag treatment. While the morphology of typical FF (A) and atypical FF (B) papillae diifer, both house K8^+^ taste buds (red). (C) After 3 days of HhAntag, the number of typical FF and atypical FF taste buds do not differ from vehicle-treated controls. (D) The number of K8^+^ pixels, a proxy for taste bud size, is comparable between vehicle and HhAntag treated mice at 3 days. (E) After 7 days, HhAntag treated mice tend to have fewer typical FF taste buds, while the number of atypical FF taste buds is significantly increased. (F) At 7 days, taste bud size does not differ between HhAntag and vehicle treated mice. Nuclei are counterstained with Draq5 (blue); dashed lines delimit the basement membrane; solid lines delimit the epithelial surface; mc = mesenchymal core. A and B are confocal compressed *z*-stacks. Scale bars = 10 μm. *N* = 3 mice for 3 days; *N* = 3-4 mice for 7 days; *n*=30 taste buds for vehicle or 30-40 for HhAntag (10 taste buds randomly selected per mouse). Data are represented as mean ± SD. Student’s t-test; \*\**P*<0.01.

### SOX2-GFP expression in FFP epithelium and taste buds is significantly decreased by inhibition of HH signaling

As taste buds are already impacted, albeit minimally, at 7 days, we reasoned that if SOX2 plays a role in taste cell renewal downstream of SHH, then SOX2 expression would be affected by short-term drug treatment. Hence, we examined GFP expression in SOX2-GFP mice treated with HhAntag or vehicle for 3 or 7 days. In intact control tongues, SOX2-GFP expression is readily detected in FFP at low magnification (Fig. 2A). At higher magnification, dimmer GFP expression was evident in FFP epithelial cells surrounding SOX2-GFP high (GFP^hi^) taste buds (Fig. 2A, inset, arrowheads and arrow respectively); this pattern of SOX2-GFP expression was similar in tongues of mice treated with drug for 3 days (Fig. 2B and inset). By contrast, in tongues from mice treated with HhAntag for 7 days, GFP expression appeared substantially reduced (Fig. 2C). Upon closer examination, GFP^hi^ signal appeared limited to apical FF taste buds (Fig. 2C, inset and arrow), with little or no GFP signal in FFP epithelium surrounding buds.

**Figure 2.**
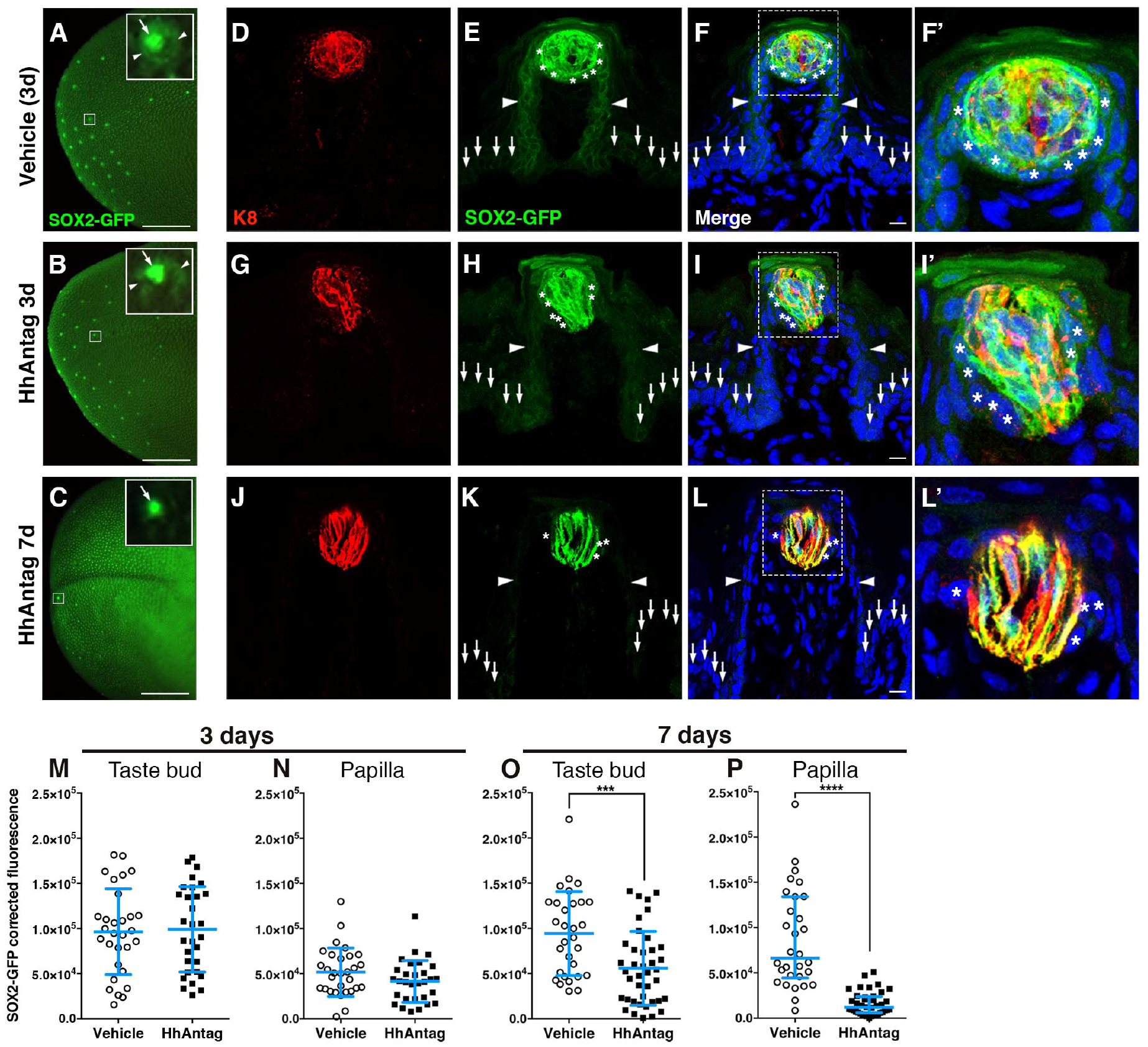
SOX2-GFP expression is reduced in taste buds and papillae in mice treated with HhAntag. (A) In whole mount preparations of vehicle treated control tongues at 3 days, taste buds appear to have high SOX2-GFP expression (GFP^hi^; insets, arrows), while adjacent FF papilla walls express lower SOX2-GFP (GFP^mid^; insets, arrowheads). (B) A similar pattern of GFP^hi^ taste buds with GFP^mid^ papilla epithelium is evident in tongues of mice treated with HhAntag for 3 days. (C) At 7 days of HhAntag treatment, fewer GFP^hi^ taste buds are detectable (low power view and inset, arrow), and papilla GFP^mid^ expression is absent. (D-L’) Immunostained tissue sections confirmed these patterns of GFP expression. (D-F’) In vehicle controls, GFP^hi^/K8^+^ cells are present in taste buds and GFP^hi^ /K8− perigemmal cells surround taste buds (asterisks); GFP^mid^ cells are restricted to FF papilla walls (arrowheads), while nearby non-taste epithelium is GFP^lo^ (vertical arrows). (G-I’) After 3 days of HhAntag, the pattern of GFP exression does not change, i.e., GFP^hi^/K8^+^ taste bud cells and K8− perigemmal cells (asterisks), GFP^mid^ papilla epithelium (arrowheads), and GFP^lo^ non-taste epithelium (vertical arrows). (J-L’) After 7 days of HhAntag, GFP^hi^/K8^+^ taste cells are reduced, GFP^hi^ perigemmal cells are lacking (asterisks) and GFP expression is virtually absent in papillary epithelium (arrowheads) and non-taste epithelium (vertical arrows). (M, N) At 3 days, SOX2-GFP corrected fluorescence intensity in taste buds plus perigemmal cells does not differ between vehicle- and HhAntag-treated mice; there is a small but non-significant decrease in GFP+ fluorescence in FFP epithelium (see methods). (O, P) After 7 days of HhAntag, SOX2-GFP intensity is significantly reduced within taste buds and perigemmal cells adjacent to buds, and within FFP walls. Nuclei are counterstained with Draq5 (blue). All images are confocal compressed *z*-stacks. A-C scale bars = 1 mm; D-L scale bars = 10 μm. *N*=3 mice 3 days; *N* =3-4 mice 7 days; *n* =30 taste buds and papillae for vehicle or 30-40 for HhAntag. Data are represented as mean ± SD, except P that are represented as median with interquartile range. Student’s t-test or Mann-Whitney U-test; \*\*\**P*<0.001, \*\*\*\**P*<0.0001.

We next examined SOX2-GFP expression in tissue sections of anterior tongues. Notably, SOX2-GFP reporter expression recapitulated SOX2 protein expression in adult tongue (see Fig 3A and Suzuki, 2008; Okubo et al., 2009; Ohmoto et al., 2017). Specifically, SOX2-GFP is highest in a subset of cells within K8+ taste buds (Fig. 2 D-F’ red), which have been proposed to represent immature and/or Type I taste cells (Suzuki, 2008). SOX2-GFP is also highly expressed (GFP^hi^) in perigemmal cells immediately surrounding each bud (Fig. 2F, F’, asterisks), more moderately (GFP^mid^) in the epithelial walls of FFP (Fig. 2 E-F arrowheads), and at low very levels in adjacent non-taste basal epithelial cells (Fig. 2F arrows). SOX2+ basal keratinocytes outside of taste buds are taste bud stem cells (Ohmoto et al., 2017), but it is not known if perigemmal SOX2hi and/or FFP wall SOX2mid cells function in this role.

**Figure 3.**
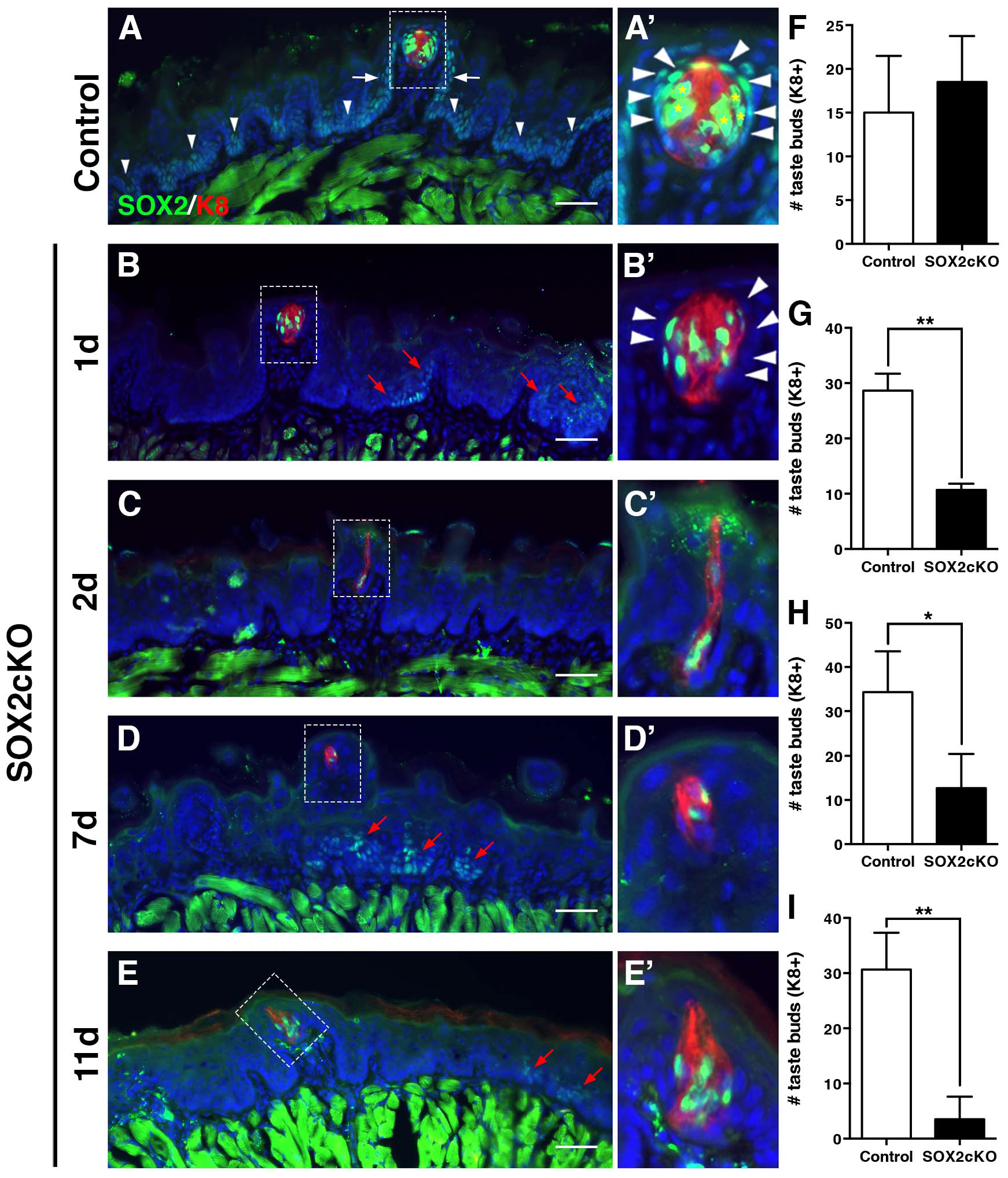
Genetic ablation of *Sox2* in K14+ progenitors disrupts taste bud renewal. (A, A’) In control mice (*K14*^+/+^*;Sox2*^*flox/flox*^), SOX2 immunoreactivity (ir)(green) is high in taste bud cells (K8-ir, red, asterisks) and perigemmal progenitor cells (arrowheads in A’). SOX2 is expressed at low levels by basal cells in papilla walls (white arrows) and non-taste epithelium (arrowheads). (B, B’) After 1 day of *Sox2* deletion (*K14*^*CreERT2/+*^*;Sox2*^*flox/flox*^), SOX2-ir cells are found within most taste buds (B’), while SOX2^+^ epithelial cells outside of buds are limited to sparse, scattered clusters (red arrows) and perigemmal cells lack SOX2 expression (arrowheads). (C-E’) A similar pattern of SOX2-ir is observed at 2 (C, C’), 7 (D, D’) and 11 (E, E’) after Cre induction; bright SOX2-immunoreactive (ir) cells are observed in occasional taste buds and scattered small clusters of more dimly SOX2-ir cells are evident in the non-taste epithelium (red arrows). (F) Taste bud number in mutant mice does not differ from controls 1 day after *Sox2* deletion (G-I) Deletion of *Sox2* results in rapid loss of K8^+^ taste buds and FF papillae. (I) K8^+^ taste bud number is dramatically and significantly reduced within 2 days of SOX2cKO. (J) A similar reduction in taste bud number is evident at 7 days post-SOX2cKO, while by 11 days, only 15% of taste buds remain in mutant tongues (K). Further, the morphology of the remaining FF taste buds is disrupted in SOX2cKO mice; taste buds have fewer and more slender taste cells (A’-E’). Nuclei are counterstained with Draq5 (blue). Scale bars=50 μm. All are fluorescence images. Scale bars = 50 μm. *N*=3-5 mice per condition. Data are represented as mean ± SD. Student’s t-test; \**P*μ0.05, \*\**P*<0.01.

GFP expression in tongues of mice treated for 3 days with HhAntag was also comparable to controls: basal cells of the FFP walls were SOX2-GFP^mid^, while perigemmal cells adjacent to taste buds were GFP^hi^ (Fig. 2G-I’, arrowheads and asterisks, respectively). K8+ taste buds (red) were also GFP^hi^ (Fig. 2G-I’), while basal progenitors in non-taste epithelium outside of FFP were SOX2-GFP^lo^) (Fig 2G-I arrows).

However, SOX2-GFP expression was significantly altered by 7 days of HhAntag. GFP was virtually absent in FFP walls (Fig. 2K, L, arrowheads), and perigemmal cells lacked GFP^hi^ expression (Fig. 2K-L’ asterisks); only elongate K8+ taste cells within taste buds remained GFP^hi^ (Fig. 2J-L’, red). These qualitative observations were confirmed by quantification of corrected SOX2-GFP fluorescence in: (1) the tissue compartment comprising the putative taste bud stem cell population, i.e., FFP walls plus perigemmal cells (Fig. 2E, H, K, arrowheads and asterisks, respectively); and (2) K8+ taste buds (Fig. 2D, G, J) (see Materials and Methods). HhAntag for 3 days did not alter SOX2-GFP expression within taste buds (Fig. 2M), and in FFP walls plus perigemmal cells, fluorescence was slightly, but not significantly reduced (Fig. 2N). However, SOX2-GFP fluorescence in both taste buds (Fig. 2O) and the FFP walls plus perigemmal compartment (Fig 2P) were significantly reduced following 7 days of HhAntag. These data indicate that HH signaling regulates SOX2 expression and suggest that HH signaling and SOX2 are required together to regulate taste bud cell renewal.

### Genetic ablation of *Sox2* in K14+ progenitors disrupts taste bud renewal

Differentiation of taste buds is perturbed in *Sox2* hypomorphic mouse embryos, indicating that SOX2 is required for early development of the taste epithelium (Okubo et al., 2006). To determine if SOX2 is required for adult taste cell renewal, we conditionally deleted *Sox2* in lingual epithelial progenitor cells by dosing *K14*^*+/CreER*^;*Sox2*^*flox/flox*^ mice once with tamoxifen (SOX2cKO). Previously, we reported that Cre expression by the *K14* allele used here results in broad although not complete activation of reporter gene expression within K14^+^ lingual progenitor cells (Castillo et al., 2014). Likewise, SOX2, as evidenced via immunostaining, was deleted in the majority of K14^+^ progenitors at 1, 2, 7 and 11 days post-tamoxifen induction (Fig 3A-E); although small clusters of SOX2*^+^* cells were scattered in the non-taste epithelium at all post-induction time points (Fig 3B, D, E, red arrows). SOX2 was also deleted from perigemmal cells, which highly express SOX2 in controls (Fig 3A’, B’ arrowheads). As taste cells do not express K14, we were not surprised to find SOX2^+^/K8^+^ taste cells in SOX2cKO tongues at all time points (Fig. 3 B’–E’), likely representing taste cells that entered taste buds and differentiated prior to *Sox2* deletion.

Despite the persistence of some taste buds, their number was dramically reduced in SOX2cKO tongues. At 1day post-tamoxifen, taste buds with normal morphology were detected in normal numbers (Fig 3B’, F). Two days post-induction, however, the number of K8^+^ taste buds was dramatically reduced in mutants compared to controls (Fig. 3G). Taste buds that remained were SOX2^+^ yet had greatly perturbed morphology; mutant taste buds were thinner and more elongated (Fig. 3C’; compare with 3A’ controls). Following this initial drastic loss, taste bud number decreased only minimally as assessed at days 7 and 11 post induction, with remaining taste buds exhibiting both distorted morphology and SOX2 protein expression (Fig. 3D’, E’, H, I), suggesting that taste cell expression of SOX2 was not sufficient to maintain taste buds.

Additionally, taste buds disappeared very rapidly in SOX2cKO mice, considering that rodent taste cells have an average lifespan of 14 days (Beidler and Smallman, 1965; Farbman, 1980). This rapid disappearance suggested that taste bud loss was not simply due to a failure of SOX2-depleted progenitors to fulfill normal replacement of taste bud cells, but rather that taste bud cells en masse were actively lost in the absence of SOX2. In control tongues, TUNEL^+^ cells are evident in the most superificial layer of the lingual epithelium, as these cells undergo cell death and form the barrier layer of the tongue (Fig. 4A). By contrast, apoptotic cells are rarely found within taste buds or basal layers of the epithelium (Fig. 4A) (Zeng and Oakley, 1999; Ichimori et al., 2009). One day after *Sox2* deletion, however, we detected many TUNEL^+^/K8^+^ taste cells in mutants compared with controls as well as increased TUNEL^+^ cells in the FFP wall plus perigemmal region (Fig. 4). By 2 days of SOX2cKO, however, TUNEL^+^ cells were only seen in superficial layers (data not shown), as in controls. Importantly, genetic deletion of *Sox2* 1 day after tamoxifen induction is restricted to basal keratinoctyes outside of taste buds, as (1) differentiated taste cells are K14^−^; (2) taste cell differentiation from adjacent stem cells requires 2-3 days following the last cell division; and (3) individual taste cells live an average of 10-14 days (see Barlow, 2015 for review). Consistent with this rationale, SOX2 expression is robust in taste bud cells at 1 day of SOX2cKO (see Fig 3B, B’), and thus, rapid taste bud loss, preceeded by a dramatic increase in taste cell apoptosis, cannot be due to SOX2 loss of function in these differentiated taste cells. Rather, we hypothesize that SOX2 acts in local taste progenitors to indirectly to support taste cell survival.

**Figure 4.**
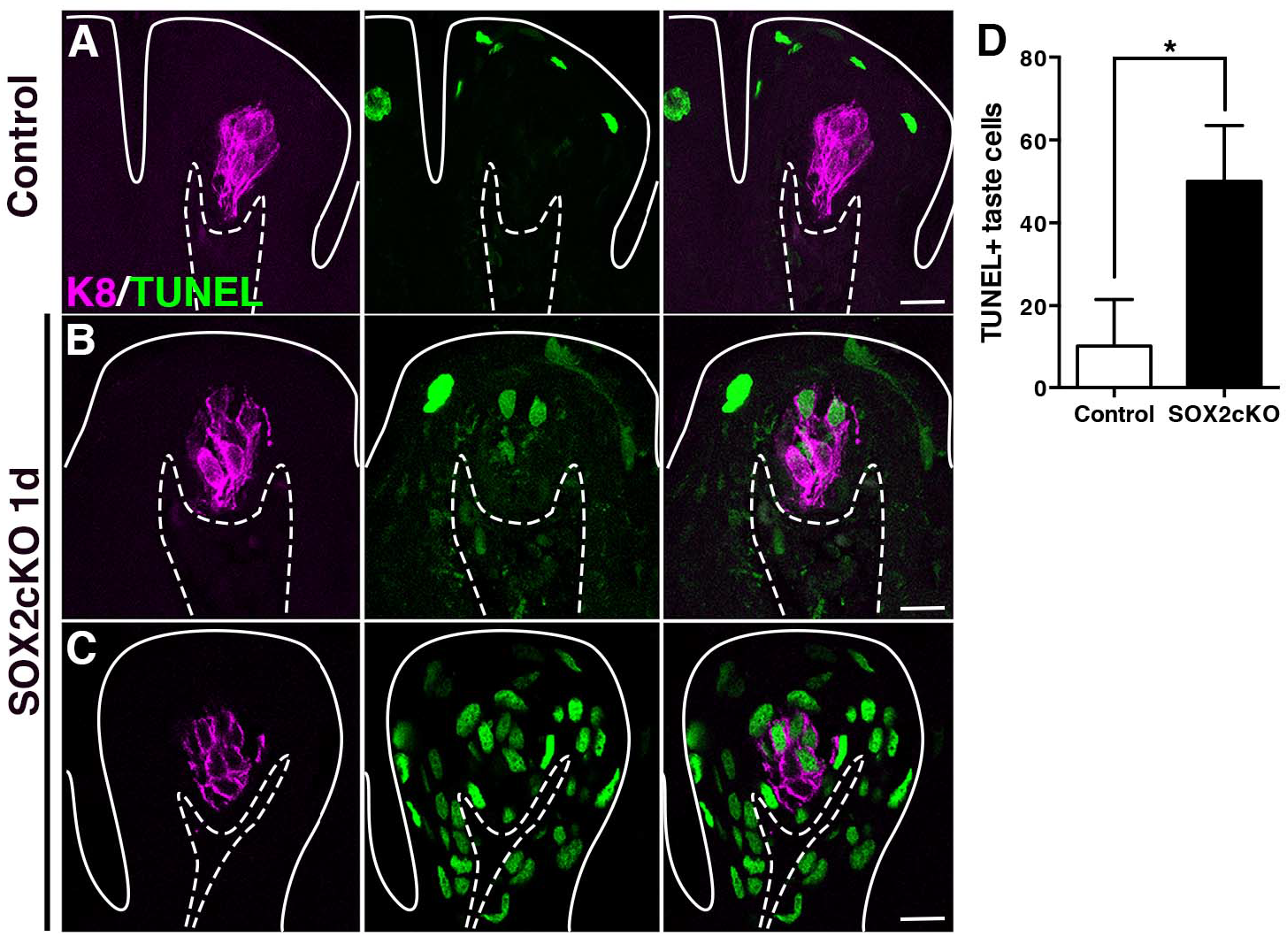
Deletion of Sox2 in progenitor cells induces taste cell apoptosis non-cell autonomously. (A) In control FFP, TUNEL+ nuclei (green) are typically detected in superficial keratinocytes as they enucleate to form the acellular surface layer of the tongue epithelium. After 1 day of SOX2cKO, TUNEL+/K8^+^ taste cells (magenta) are detected in numerous mutant FFP (B), and occasional FFP are found to have extensive TUNEL+ cells (C). Dashed lines delimit the basement membrane; solid lines delimit the epithelial surface. Scale bars = 20 μm. (D) At 1 day after SOX2cKO, mutant mice have significantly more TUNEL+/K8^+^ taste cells than controls. *N*=3 mice per condition. Data are represented as mean ± SD. Student’s t-test; \**P*<0.05.

### Loss of SOX2 in K14^+^ progenitors blocks differentiation of taste and non-taste epithelial cells

In addition to increased taste cell death, we reasoned that rapid loss of taste buds in the absence of SOX2 might be due to altered cell fate specification within the progenitor population. Hence, we next investigated if SOX2cKO affected differentiation of K14^+^ progenitors. In control taste papillae, K14^+^ basal progenitors are found lining the FFP walls and perigemmaly (Fig. 5A). The pattern of K14 expression is similar 1 day after SOX2cKO (data not shown). By contrast, at 2 days post-induction, in SOX2cKO tongues K14^+^ cells expand above the basal epithelial layer and encroach upon FF taste buds (Fig. 5B, arrowheads). By 7 and 11 days after Cre induction, many more K14^+^ cells occupy the suprabasal epithelium, including regions formerly occupied by taste buds (Fig. 5C, D). Additionally, at later time points, suprabasal K14^+^ cells were elongated and flattened, in contrast to basally located, ovoid K14^+^ cells (Fig 5C, D
, red arrows).

**Figure 5.**
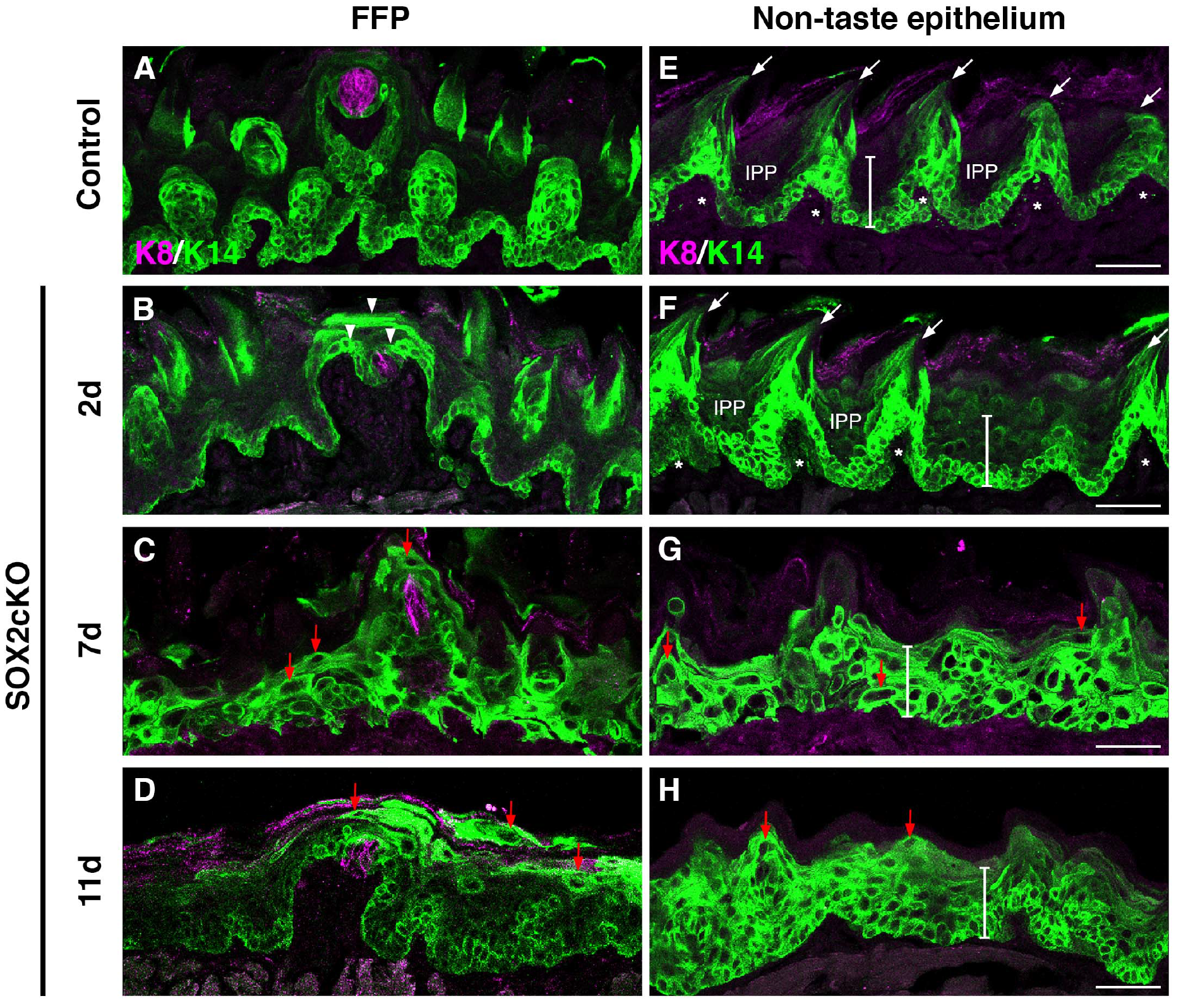
Loss of SOX2 in K14+ progenitors blocks fate acquisition of taste and non-taste epithelial cells. (A) In control FFP, K14^+^ progenitors (green) are limited to the basal epithelial layer, and adjacent to taste buds (K8, magenta). (B) By 2 days of Sox2 deletion in *K14*^*CreERT2/+*^;*Sox2*^*flox/flox*^ (SOX2cKO) mice, K14^+^ cells reside in suprabasal epithelial layers and have expanded around K8^+^ taste buds (arrowheads). (C, D) By 7 and 11 days post-SOX2cKO, K14^+^ cells comprise most of the FF epithelium, and many of these cells have enlarged cell somata with elongated K14^+^ processes (red arrows). (E) Control non-taste epithelium is characterized by basally located K14^+^ progenitors and filiform papillae (arrows, asterisks mark each filiform papilla core), interspersed by K14^−^ interpapillary (IPP) regions. The white bar spans the epithelium from the basement membrane to the superficial cellular layers (73 μm). (F) Filiform papillae are initially evident at 2 days of SOX2cKO, but K14^+^ cells are uncharacteristically detected in suprabasal layers. At 7 (G) and 11 (H) days of SOX2cKO, filiform papillae are no longer evident, K14^+^ cells span the entire thickness of the epithelium (white bars), and these K14^+^ cells have atypical morphologies (red arrows). Scale bars = 50μm.

As SOX2 is expressed in non-taste epithelial progenitors, we reasoned that in the absence of SOX2, keratinocyte differentiation might also be disturbed throughout the tongue. Mechanosensory filiform papillae with small, short pits between them make up the majority of the non-taste epithelium. K14^+^ progenitors in control tongues surround the mesenchymal core of each filiform papilla (Fig. 5E, asterisks), occur in filiform papilla *per se* (Fig. 5E, arrows), and lie at the basement membrane of the interpapillary pits (Fig. 5E, “IPPs”); this pattern did not differ 1 day after SOX2cKO (data not shown.) At 2 days following *Sox2* deletion, spinous-shaped filiform papillae with mesenchymal cores were present (Fig. 5F, arrows and asterisks), however, IPPs were now comprised mostly of K14^+^ progenitors instead of K14^−^ differentiated keratinocytes, and layers of K14^+^ cells within the IPPs extended to the tongue surface (Fig. 5I, IPP and white vertical bars). At 7 and at 11 days post-induction, filiform papillae were lost altogether and K14^+^ progenitor cells expanded throughout the entire depth of the non-taste epithelium (Fig. 5G, H, white vertical bars).

To determine if K14^+^ progenitors expand in the lingual epithelium because of sustained proliferation, we used Ki67 to detect actively cycling cells in tissue sections of the anterior tongue (Fig. 6A, A’). The majority of progenitor cells are Ki67^+^ in control mice, and these proliferating cells are restricted to the basal layer (Fig. 6B, B’). At 2 days of SOX2cKO, proliferating cells appeared disorganized, with numerous Ki67^+^ cells in suprabasal layers of the epithelium (Fig 6C, C’, arrowheads). With prolonged SOX2 loss, progressively more proliferating cells were detected both basally and suprabasally (Fig. 6D-E’, arrows (basal cells) and arrowheads (suprabasal cells)).

**Figure 6.**
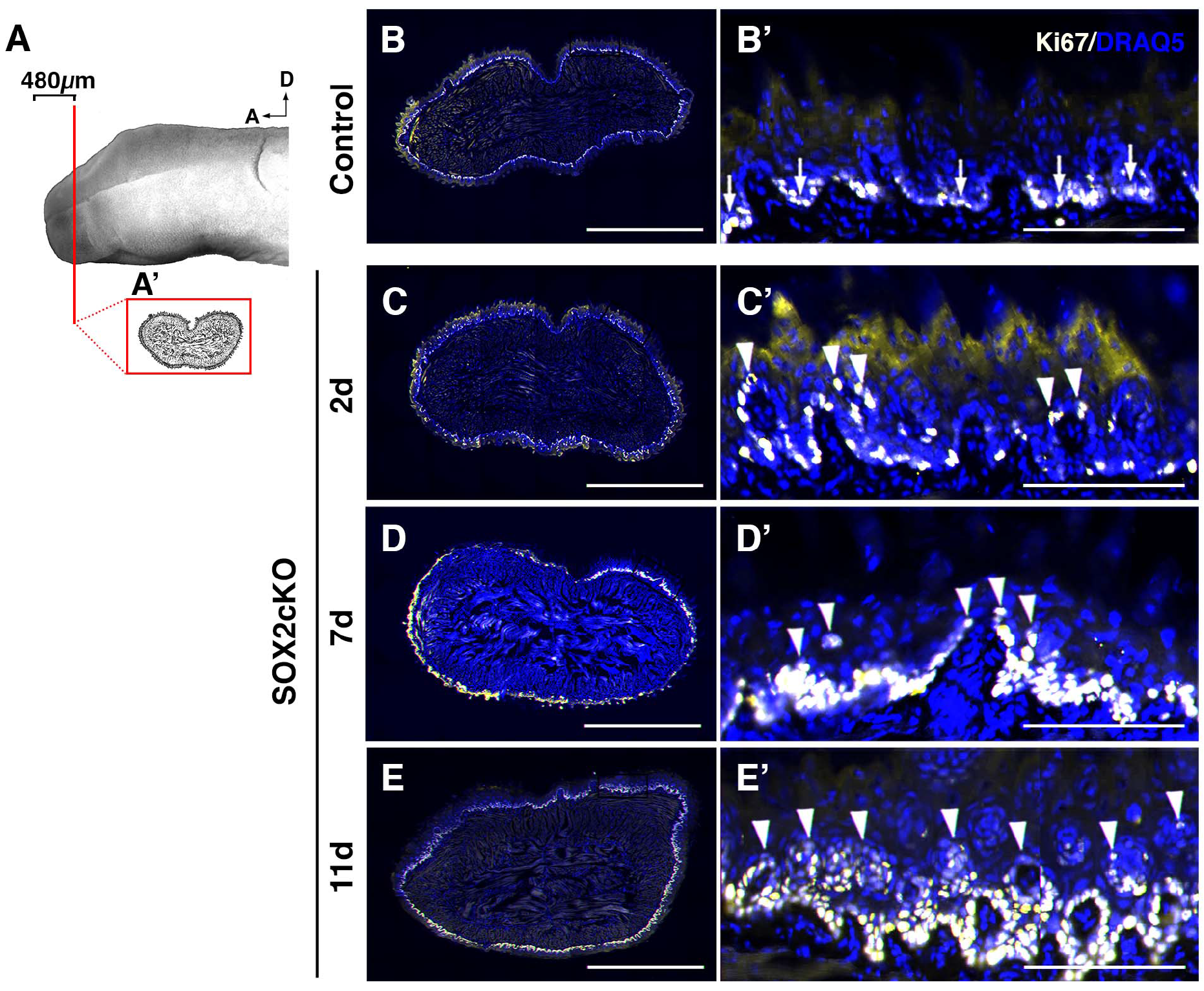
Proliferation is disorganized in SOX2cKO mice. (A) Proliferating cells (Ki67^+^) were quantified in representative transverse sections through the anterior tongue (A’) (480 μm from the tip). (B, B’) In controls, Ki67^+^ (light yellow) proliferating cells are restricted to the basal layer of the lingual epithelium. (C-E’) In SOX2cKO mice, Ki67+ cells are also found basally, but progressively more Ki67^+^ cells are found in suprabasal layers at later times post SOX2cKO. (C’-E’ arrowheads). Nuclei are counterstained with Draq5 (blue). All images are scanned best focus sections. B-E scale bars = 1 mm; B’-E’ scale bars = 125 μm.

Thus, our data indicate that genetic deletion of *Sox2* in lingual progenitor cells impairs proper taste and non-taste cell fate specification, promotes overexpansion of progenitor cells via aberrant proliferation in non-taste epithelium, and activates cell death in taste epithelium. The combination of these cellular events likely underlies the swift decrease in FF taste buds and loss of filiform papillae.

### Overexpression of Shh in SOX2cKO lingual progenitors transforms Shh from a pro-taste differentiation factor to an epithelial mitogen

SHH over-expression in K14^+^ progenitors cells induces *de novo* differentiation of taste buds in regions of the lingual epithelium formerly thought incapable of sustaining taste cell differentiation (Castillo et al., 2014). Hence, we next asked whether Shh over-expression in K14^+^ progenitors can drive taste cell differentiation in the absence of SOX2. We took advantage of a *Shh* conditional knock-in allele, SHH-IRES-YFPcKI (SHHcKI), together with SOX2cKO to drive SHH expression and *Sox2* deletion in K14^+^ progenitors of adult *K14*^*+/CreER*^*;Rosa*^*Shh-IRES-YFPcKI*^*;Sox2*^*flox/flox*^ mice (SHHcKI-SOX2cKO). Control and mutant mice were given a single tamoxifen dose and tongues analyzed at 11 days. In genetic controls (*Rosa*^*Shh-IRES-YFPcKI*^*;Sox2*^*flox/flox*^ treated with tamoxifen), FF taste buds are readily visible on the tongue surface as translucent ovoids (Fig. 7A, arrowheads) interspersed among spinous filiform papillae. In SHHcKI-SOX2cKO mice, neither FF nor filiform papillae were abundant on the tongue surface; those FFP that remained likely reflected mosaic Cre activation (Fig. 7B, arrowheads). The loss of taste buds was confirmed in K8-immunostained tissue sections; SHHcKI-SOX2cKO mice exhibited on average one K8^+^ FF taste bud in the anteriormost 1.5 mm of the tongue per mouse compared to ~20 buds evident in the same region of each tongue in controls (Fig. 7C). Thus, deletion of *Sox2* prevents SHH-dependent ectopic taste bud formation, a result consistent with our previous observation that ectopically-induced taste buds expressed high levels of SOX2 (Castillo et al., 2014).

**Figure 7.**
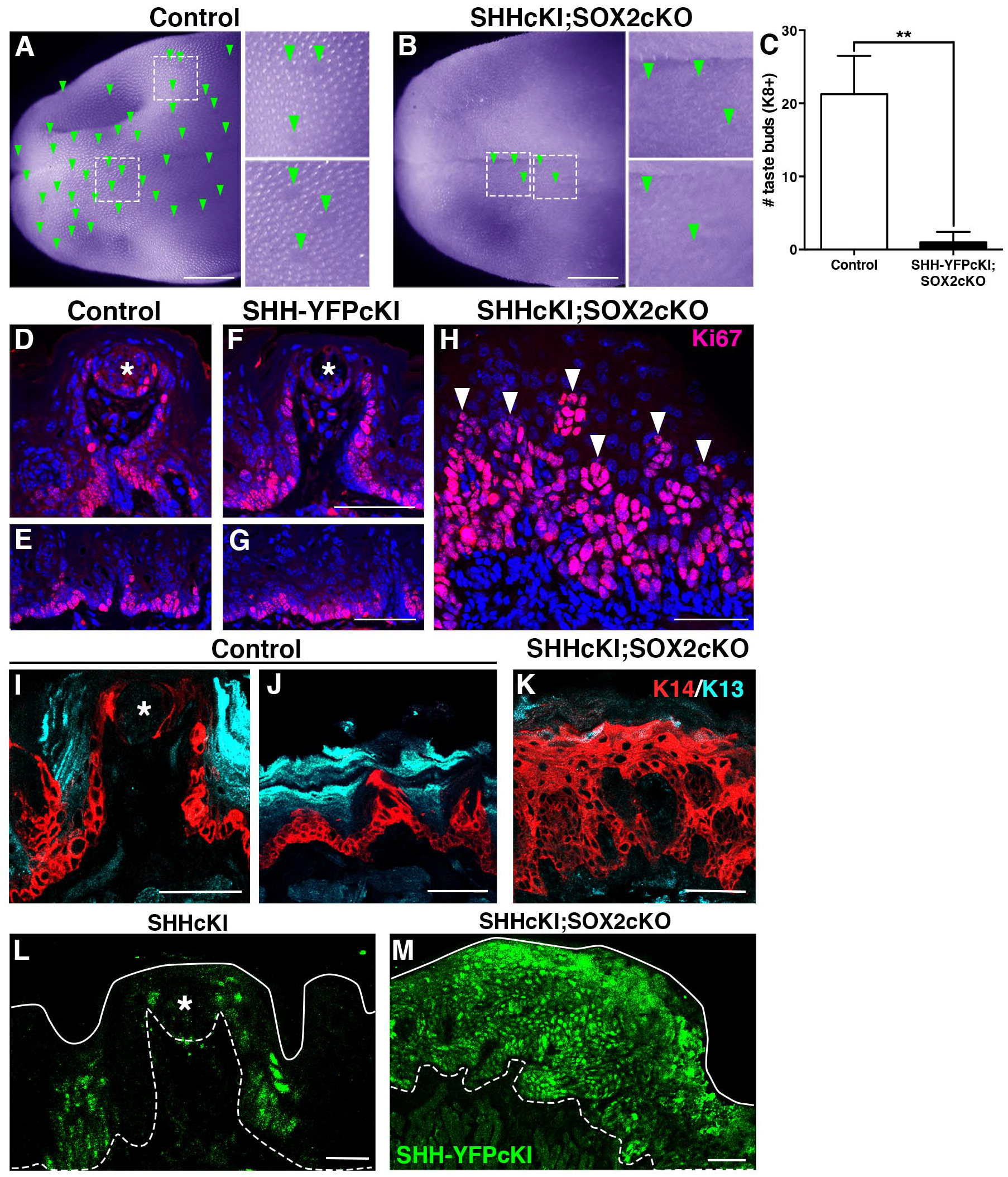
Overexpression of SHH in SOX2cKO lingual progenitors transforms SHH from a pro-taste differentiation factor to an epithelial mitogen. (A) In a control tongue (*Rosa*^*Shh-IRES-YFPcKI*^;*Sox2*^*flox/flox*^) viewed in whole mount (pseudocolored purple to enhance contrast), FF papillae are evident as clear ovals (green arrowheads, insets) embedded within the spinous filiform papillae that cover the tongue surface. (B) In double mutant tongues (SHHcKI;SOX2cKO in K14^+^ progenitors) at 11 days, FFP (green arrowheads and insets) and filiform papillae are mostly absent in tongues. (C) Tallies of taste buds in immunostained tissue sections from the first 1.5 mm of the tongue show K8^+^ taste buds are drastically diminished in SHHcKI;SOX2cKO mice compared with the same region in controls. (D-H) In control and SHHcKI tongues, actively cycling cells (Ki67, red) have the same basal distribution as K14^+^ progenitors in taste (D, F; taste bud, asterisk) and non-taste (E, G) epithelia. (H) In the absence of SOX2, SHH over-expression massively increases epithelial proliferation (Ki67^+^ red, arrowheads). (I) In control mice, K14^+^ progenitors (red) are present adjacent to taste buds (asterisk) and in FFP walls, and K13^+^ (cyan) differentiated keratinocytes make up the surface of the FFP. (J) In non-taste epithelium, K14^+^ progenitors reside basally, while differentiated K13^+^ keratinocytes are found suprabasally. (K) Lingual epithelium of SHHcKI;SOX2cKO mice is populated almost exclusively by K14^+^ cells at the expense of differentiated K13^+^ cells. (L) SHH-YFPcKI^+^ patches (green) 14 days post-Cre induction are limited to discrete patches in the presence of SOX2, as reported previously (Catillo et al. 2014). (M) In the absence of SOX2 11 days post-Cre induction, SHH-YFPcKI+ patches (green) are significantly expanded. Nuclei are counterstained with Draq5 (blue); dashed lines delimit the basement membrane; solid lines delimit the epithelial surface. D-M are compressed confocal *z*-stacks. A and B scale bars = 1 mm; D-K and M scale bars = 50 μm; L scale bar = 20 μm. *N*=4 mice per condition. Data are represented as mean ± SD. Student’s t-test; \*\**P*<0.01.

We also assessed the gross morphology of the dorsal, lateral and ventral epithelia in tongue cryosections from mutants and controls. In general, despite induction of ectopic taste buds, lingual epithelium is normally structured in SHHcKI mutants. In control and SHHcKI mice, FFP, readily identifiable via nuclear counterstain, are found predominantly on the dorsal surface of the tongue (Fig. S1A, B, arrowheads), while in SHHcKI-SOX2cKO mutants, FF taste buds were mostly absent (Fig. S1C). Dorsal, lateral, and ventral epithelia in control and SHHcKI tongues have well organized fungfiform and filiform papillae with discrete mesenchymal cores (Fig. S1D, E, G, H, asterisks), and there is a clear distinction between the basal epithelial layer and the lamina propria, i.e., connective tissue, below (Fig. S1D, E, G, H). In SHHcKI-SOX2cKO mice, however, papillary epithelial invaginations with mesenchymal cores virtually disappear, the epithelium appears to contain more cells, and the boundary between epithelium and lamina propria is indistinct (Fig. S1C, F, I). This disorganized phenotype is more severe in the lateral and ventral aspects of the tongue, where, in controls, taste buds do not typically reside.

We next examined the extent of epithelial proliferation in control, SHHcKI, and SHHcKI-SOX2cKO tongues, reasoning that an increase in proliferation may underlie the aberrant morphology seen in SHHcKI-SOX2cKO lingual epithelia. In both controls and SHHcKI mice, Ki67^+^ cells were restricted to the basal epithelial layer of both FFP and non-taste epithelium, and the proportion of proliferating epithelial cells appeared comparable in control and SHHcKI tongues (Fig. 7D-G); these observations are consistent with the minimal impact of loss of SHH signaling on lingual epithelial proliferation (Castillo-Azofeifa et al., 2017; Kumari et al., 2017). By contrast, but similar to the impact of SOX2cKO alone (see Fig. 6E’), Ki67^+^ cells were dramatically increased and found throughout a greatly expanded epithelium in SHHcKI-SOX2cKO tongues (Fig. 7H). The expanded domains in double mutant epithelium comprised primarily K14^+^ progenitors, lacked differentiated K13^+^ keratinocytes that occupy suprabasal layers in controls (Fig. 7I-K), and coincided with large areas of SHH-overexpressing cells (Fig 7L, M).

## Discussion

Little is known about the functional role of SOX2 in adult lingual epithelium, as well as the connection, if any, between HH signaling and SOX2. Specifically, we wanted to investigate if the interaction of these two factors is crucial for taste bud homeostasis. Previously, we showed that SHH overexpression in K14^+^ progenitors of the adult non-taste epithelium results in elevated SOX2 levels, and these ectopic patches of high SHH/SOX2 expression coincide with the development of ectopic taste buds (Castillo et al., 2014). These findings suggested that the mechanism by which SHH regulates taste bud homeostasis is by increasing SOX2 expression levels.

In the present study, we show that HH signaling inhibition results in rapid downregulation of SOX2 in both taste progenitors and intragemmal taste bud cells. Additionally, *Sox2* conditional deletion in lingual progenitor cells promotes taste cell death, prevents differentiation of taste cells and lingual keratinocytes, and expands the undifferentiated progenitor cell population. Finally, we demonstrate that SHH functions as a mitogen rather than a pro-taste bud differentiation factor in the absence of SOX2. Together, these results demonstrate that SOX2 is downstream of the HH signaling cascade and that SOX2 is essential for translating HH activity into the appropriate cellular output.

We first sought to elucidate if acute use of an HPI impacts taste bud maintenance, as chronic administration of HPIs in humans and mice affects the taste system (LoRusso et al., 2011; Tang et al., 2012; Rodon et al., 2014; Kumari et al., 2015; Yang et al., 2015; Kumari et al., 2017). After one week of HhAntag treatment, taste bud number and size trended downward, while the incidence of degenerating FFP was significantly increased. Even though these differences were small or not statistically significant, they nonetheless suggested that HH dependent molecular mechanisms regulating taste cell renewal were already affected by 7 days but had not yet been translated into a robust cellular phenotype.

In the adult tongue, SOX2 is highly expressed in perigemmal cells and in a subset of intragemmal taste bud cells, whereas SOX2 expression is low in the basal keratinocytes of the non-taste epithelium (Okubo et al., 2006; Suzuki, 2008; Okubo et al., 2009; Ohmoto et al., 2017). In adult mice, after transection of the glossopharyngeal nerve innervating the posterior circumvallate taste papilla, taste buds disappear, as does SOX2 expression in taste progenitors and taste buds. Nerve regeneration is required for taste bud regeneration (Cheal and Oakley, 1977) and similarly, nerve regeneration leads to reappearance of high levels of SOX2 expression in perigemmal cells followed by expression within the regenerating taste buds (Suzuki, 2008), suggesting SOX2 is involved in adult taste bud regeneration.

Monitoring SOX2 activity via SOX2-GFP expression in HhAntag treated mice, we found that SOX2 expression was significantly decreased in FFP taste epithelium after 7 days of drug treatment. Interestingly, with HPI treatment, we detected a greater effect on SOX2-GFP expression in the perigemmal progenitors compared to SOX2-GFP inside taste buds, which aligns with the perigemmal pattern of recovery of SOX2 expression prior to taste bud regeneration after denervation (Suzuki, 2008). In sum, these data support the idea that blocking HH signaling: (1) triggers downregulation of SOX2 expression in HH-responding perigemmal progenitors, followed by (2) loss of SOX2 expression in mature taste cells, and finally (3) taste bud regression.

Prior studies have suggested that SOX2 is required but not sufficient for embryonic taste bud formation. *Sox2* hypomorphic embryos, which express ~20% of normal SOX2 levels, fail to form differentiated taste buds at birth, while *Sox2* gain-of-function does not induce taste bud differentiation (Okubo et al., 2006). Our findings extend a requirement for SOX2 to adult taste bud renewal, as loss of SOX2 in adult tongue progenitors causes taste bud deterioration. Compared to the effect of HH signaling inhibition, however, *Sox2* deletion prompts an extremely rapid decline in taste epithelium, with taste bud regression already evident within 2 days of conditional deletion. Thus, in addition to the well recognized trophic role of the gustatory innervation, progenitor cells likely provide pro-taste bud survival factors downstream of SOX2.

A comprehensive study showed that SOX2 is expressed in many adult mouse epithelial tissues (e.g. tongue, lungs, lens, glandular stomach, esophagus, forestomach, and anus), where it marks basally situated progenitor cells (Arnold et al., 2011). This group showed that ablation of SOX2^+^ basal cells in the tongue and oral mucosa resulted in inflammation, ulcers and edema in the oral cavity (Arnold et al., 2011). Our data complement this study, as loss of SOX2 in tongue progenitors alters adult lingual epithelium homeostasis by promoting K14^+^ progenitor proliferation and impeding taste and non-taste cell fates. However, our results contrast with reports that hypomorphic SOX2 expression did not affect differentiation of filiform papillae and progenitor proliferation in embryos (Okubo et al., 2006), as well as that stratified epithelial layers in the tongue are unaltered by ablation of SOX2^+^ progenitors in adults (Arnold et al., 2011). The discrepancies with the developmental study may be due to a more limited function of SOX2 in embryonic lingual epithelium, as opposed to the broader role for SOX2 in adult tongue, where we show it is required for both taste and non-taste epithelium. Alternatively, hypomorphic SOX2 expression may be sufficient for filiform papillae development, where in adults SOX2 levels are lowest. Our findings, however, parallel those from studies of SOX2^+^ dental epithelial cells that contribute to all epithelial cell lineages of the mouse incisor (Juuri et al., 2012).

Our data support a model in which HH controls tongue epithelium homeostasis by regulating SOX2 levels in lingual progenitors. In support of this hypothesis, in adult FFP, the perigemmal cells that highly express SOX2 are also GLI1^+^, and are directly affected by LDE225, an HPI (Liu et al., 2013; Kumari et al., 2015). Furthermore, a SOX2-SHH link has been reported in embryonic tongues, where both genes are coexpressed in placodes of the developing taste papillae, and thus SOX2 and SHH may also interact in taste bud development. Imbalance between several signaling circuits including SHH and SOX2 causes elevated levels of SOX2 in cells that normally differentiate into keratinocytes, and in some cases these cells appear to form taste buds (Beites et al., 2009). A direct relationship between SHH and *Sox2* has been documented in telencephalic neuroepithelial cells, where SOX2 expression is under the control of the GLI2 transcription factor (Takanaga et al., 2009). Further, in non-small cell lung cancer stem cells, GLI1 was found to bind to the promoter region and regulate *Sox2* transcription (Bora-Singhal et al., 2015).

To test the requirement of *Sox2* downstream of HH in inducing taste bud differentiation, we overexpressed SHH in lingual epithelial progenitors and simultaneously deleted *Sox2*. We found that instead of inducing ectopic taste buds, SHH activated hyperproliferation of K14^+^ basal progenitors and blocked taste and non-taste epithelial differentiation. The tongue epithelial architecture was profoundly changed, developing into a hyperplastic basal cell carcinoma-like phenotype (Oro et al., 1997; Kasper et al., 2012; Wong and Dlugosz, 2014). SHH has been shown to function as a mitogen in a variety of tissues under homeostasis. In mouse lung development, SHH regulates cell proliferation of the epithelium and mesenchyme (Bellusci et al., 1997). In the anagen hair follicle, transit amplifying progeny signal via SHH to the quiescent stem cell pool to proliferate for continual hair regeneration (Hsu et al., 2014), and conditional SHH signaling activation in the adult brain results in expansion of neural stem cells at the expense of their progeny (Ferent et al., 2014). Moreover, several types of cancers are associated with mutations of the HH pathway, including basal cell carcinomas (Rubin and de Sauvage, 2006; Jiang and Hui, 2008; Ng and Curran, 2011; Petrova and Joyner, 2014), and in a mouse model, SHH overexpression in K14^+^ cells in the skin causes formation of basal cell carcinoma (Oro et al., 1997). Interestingly, in the tongue epithelium, SHH over-expression alone induces taste bud differentiation; only in the absence of SOX2 does SHH expression cause massive hyperproliferation and basal cell expansion, characteristic of basal cell carcinomas. Interestingly, while SOX2 expression is amplified in many cancers (e.g. Boumahdi et al., 2014), SOX2 expression can be protective in gastric tumors (Sarkar et al., 2016) and oral squamous cell carcinoma (Fu et al., 2016), suggesting SOX2 loss in the tongue may be pro-oncogenic.

In summary, our results demonstrate that in adult tongue, HH signaling functions through SOX2, and SOX2 is required for maintenance and renewal of both taste buds and non-taste lingual epithelium. Overall, our findings suggest SOX2 is a molecular gatekeeper of HH signaling and possibly other signaling pathways in the adult tongue. Going forward, determining whether *Sox2* is a direct or indirect downstream target of the HH signaling will help develop therapies for mitigating taste disruption due to the use of HPIs as chemotherapy and advance our understanding of the molecular mechanisms of lingual epithelium homeostasis.

## Materials and methods

### Animals

Male and female mice were all on a mixed background. Mouse lines used include combinations of the following alleles or transgenes: *Sox2*^*GFP*^(Arnold et al., 2011), *K14*^*CreER*^ (Li et al., 2000), *Sox2*^*flox*^ (Shaham et al., 2009), *R26R*^*Shh-IRES-YFPcKI*^ (Castillo et al., 2014). Mice were 6-12 weeks of age at the start of each experiment and data for this study were gathered from at least 3 mice per time point. Mice were genotyped as previously described (Shaham et al., 2009; Arnold et al., 2011; Castillo et al., 2014) and rodent work was done in accordance to approved protocols by the Institutional Animal Care and Use Committee at the University of Colorado Anschutz Medical Campus and University of California San Francisco.

### HhAntag administration

HhAntag was prepared as described by Yauch et al. (2008) and was administered to *Sox2*^*GFP*^ mice via oral gavage twice daily at a dose of 100 mg/kg for 3 or 7 days.

### Tamoxifen induction of Cre

To delete Sox2 in lingual epithelial cells by Cre activation, *K14*^*CreER*^;*Sox2*^*flox/flox*^ mice were gavaged once with a dose of 5 mg tamoxifen (T5648, Sigma) dissolved in corn oil; mice were sacrificed 1, 2, 7 or 11 days from the start of the experiment. To misexpress Shh and delete Sox2 in lingual epithelial, *K14*^*CreER*^;*R26R*^*Shh-IRES-YFPcKI*^;*Sox2*^*flox/flox*^ mice received a single tamoxifen dose of 5 mg and tissue was collected 11 days post-tamoxifen induction.

### Tissue preparation

Harvested tongues were fixed by immersion or perfusion. Immersion-fixation: animals were euthanized by CO2 inhalation followed by cervical dislocation. Tongues were dissected, rinsed in sterile ice-cold 1x Phosphate Buffered Saline (PBS), and immersed in 4% paraformaldehyde (PFA) in 0.1 M PB overnight at 4°C. Tissue was embedded in Tissue-Tek^®^ O.C.T.^™^ Compound (4583, Sakura), frozen, and stored at −80°C. Perfusion-fixation: animals were anesthetized by i.p. injection of 250 mg/kg Avertin (2,2,2-Tribromoethanol) and transcardially perfused with Periodate-Lysine-Paraformaldehyde (PLP) (Pieri et al., 2002). Dissected tongues were post-fixed in PLP for 3 hours at 4°C and then cryoprotected in 20% sucrose in 1x Phosphate Buffer (PB) overnight at 4°C. Tissue was embedded in Tissue-Tek^®^ O.C.T.^™^ Compound, frozen, and stored at −80°C. Processing of immersion or perfusion-fixed tongues was restricted to the anterior 1.5 mm of the tongue with a high density of fungiform papillae. Eight sets of serial cryosections (12 μm) per tongue were collected on Superfrost Plus Slides (12-550-15, Fisher Scientific).

### Immunofluorescence

Immunofluorescence was performed on immersion or perfusion-fixed 12 μm cryosections as described (Nguyen and Barlow, 2010). Primary antisera and dilutions: rat anti-K8 (Troma) (1:250; Developmental Studies Hybridoma Bank, University of Iowa), chicken anti-GFP used to detect GFP or YFP (1:1000; GFP-1020, Aves Labs), goat anti-SOX2 (1:500; sc-17320, Santa Cruz), rabbit anti-K14 (1:3500; PRB-155P, Covance), rabbit anti-Ki67 (1:200; RM-9106-S, Thermo Fisher Scientific) and guinea pig anti-K13 (1:500; BP5076, Acris Antibodies). Appropriate secondary antisera from Thermo Fisher Scientific (A11006, A11081, A21247, A21208, A11039, A11055, A11008, A11010, A21245, A21206, A31573, S11225, A11073), Jackson ImmunoResearch (712-165-153, 712-605-150) and Vector Laboratories (PK-6101) were used at 1:1000 (host: goat), 1:800 (host: donkey), and 1:500 (rabbit IgG biotinylated). Sections were counterstained with Draq5 (1:8000; 108410, AbCam), Sytox Green Nucleic Acid Stain (1:50000, S7020, Thermo Fisher Scientific) or Dapi (1:10000; D3571, Thermo Fisher Scientific), and coverslipped with Fluormount G (0100-01, SouthernBiotech) or ProLong Gold Antifade (P36930, Thermo Fisher Scientific).

TUNEL assay was performed on immersion-fixed 12 μm cryosections as described (Gaillard et al., 2015) using In Situ Cell Death Detection Kit, TMR red (12156792910, Roche).

### Image acquisition and analysis

All image acquisition and analyses were performed blind to condition. Fluorescence and bright-field images were acquired using a Zeiss Axioplan II microscope, an Olympus SZX12 stereo microscope or Leica DM5000 B, a Retiga 4000R camera with Q-Capture Pro-7 software, an Axiocam CCD camera with Axiovision software or Leica DFC 500 with LAS V4.9 software. Confocal images were obtained as a *z*-stack of 0.76 μm optical sections acquired sequentially using a Leica TCS SP5 II confocal microscope with LASAF software or Zeiss Oberver Z1 with ZEN blue software. Whole tissue section scannings were acquired sequentially using a Leica DFC 365FX camera on a Leica DM6000B microscope with the imaging software Surveyor by Objective Imaging. A series of 20x images was obtained for each flourophore (Texas Red, Fitc, Cy-5), aligned and stitched together using the Best Focus option in the Surveyor software. The final rendering is a mosaic RGB image of each section.

The most anterior 1.5 mm of each tongue was collected as 8 serial sets of 16 cryosections and a single series was used for each of the different immunomarkers. Each fungiform taste bud was counted if: (1) it was found within a FF papilla; and (2) it housed at least 1 K8-immunoreactive (K8^+^) cell with a nuclear profile. Additionally, fungiform taste buds were categorized as follows (and see Fig. 1A, B): (1) Typical Fungiform Papilla and taste bud (Typical FFP): papilla with epithelial invagination into the lamina propria mesenchyme, where a mesenchymal core is defined by a basal epithelial layer; the papilla apex has a plateau-like surface housing a single taste bud. Each taste bud has a characteristic onion-like shape and is composed of fusiform cells. (2) Atypical Fungiform Papilla and taste bud (Atypical FFP): papilla and mesenchymal core are narrow; the papilla apex is filiform and houses a single taste bud. The taste bud is narrow and has fewer taste cells; the remaining taste cells have a stretched appearance (Oakley et al., 1990; Nagato et al., 1995). All taste buds in either type of papillae were tallied and assigned a number. Typical and atypical FF taste buds were analyzed separately. Sets of 10 typical FF taste buds per mouse were randomly selected (random.org) for quantification of total number of K8^+^ pixels inside taste buds. We analyzed each confocal optical section from every taste bud *z*-stack using our imstack toolbox developed in MATLAB (Mathworks, Natick, MA) (Castillo-Azofeifa et al., 2017). We loaded each *z*-stack into the imstack toolbox and established a rectangular region-of-interest (ROI) that completely encompassed the taste bud. The same ROI dimensions were used for all analyzed images. Signal in all 3 channels was thresholded using Otsu’s method (Otsu, 1979). Signal was designated as only those pixels with an intensity above the calculated threshold value within the taste bud ROI.

We measured the corrected integrated density (CID) of SOX2-GFP immunofluorescence from *z*-stacks using the open-source platform Fiji (Schindelin et al., 2012). We first established a taste bud region-of-interest (ROI^a^) encompassing each K8^+^ taste bud (red). SOX2-GFP CID was quantified within the ROI^a^ to obtain SOX2-GFP corrected fluorescence within each taste bud. To quantify SOX2-GFP corrected fluorescence of FFP walls plus perigemmal cells, we set a new ROI (ROI^b^) delimiting the papilla walls and taste bud. However, in ROI^b^ we masked the area corresponding to the taste bud ROI^a^, and quantified SOX2-GFP CID within the ROI^b−a^ to obtained SOX2-GFP corrected fluorescence within each papilla. CID was obtained using the following calculation: Corrected integrated density = Integrated density – (Area selected × Mean value of background) (Gavet and Pines, 2010).

For whole section analysis of cell proliferation by Ki67 immunoreactivity, representative sections were selected from a region of the tongue where fungiform and filliform papillae are adequately distributed for the analysis. This region of the tongue is located 480 μm from the tip of the tongue.

### Statistical analysis

Normally distributed data were analyzed using the parametric two-tailed Student’s t-test with Welch’s correction. The non-parametric Mann–Whitney U-test was used if the data did not fit a normal distribution. Significance was taken as *P*<0.05 with a confidence interval of 95%. Data are presented as mean ± SD for parametric data or as median with interquartile range for non-parametric data.

## Acknowledgements

We thank Kelly Zaccone, Dong-Kha Tran, Sarah Alto, Rebecca d’Urso and Pauline Marangoni for excellent mouse assistance; and Fred deSauvage and Genentech for HhAntag.

## Author contributions

Conceptualization: D.C.-A., K.S., O.D.K., L.A.B.; Methodology: D.C.-A., K.S., O.D.K., L.A.B.; Formal analysis: D.C.-A., L.G., B.J.; Investigation: D.C.-A., K.S., O.D.K., L.A.B.; Writing - original draft: D.C.-A., L.A.B.; Writing - review & editing: D.C.-A., K.S., O.D.K., L.G., L.A.B.; Visualization: D.C.-A.; Supervision: L.A.B.; Project administration: L.A.B.; Funding acquisition: O.D.K. and L.A.B.

---

**Figure S1.**
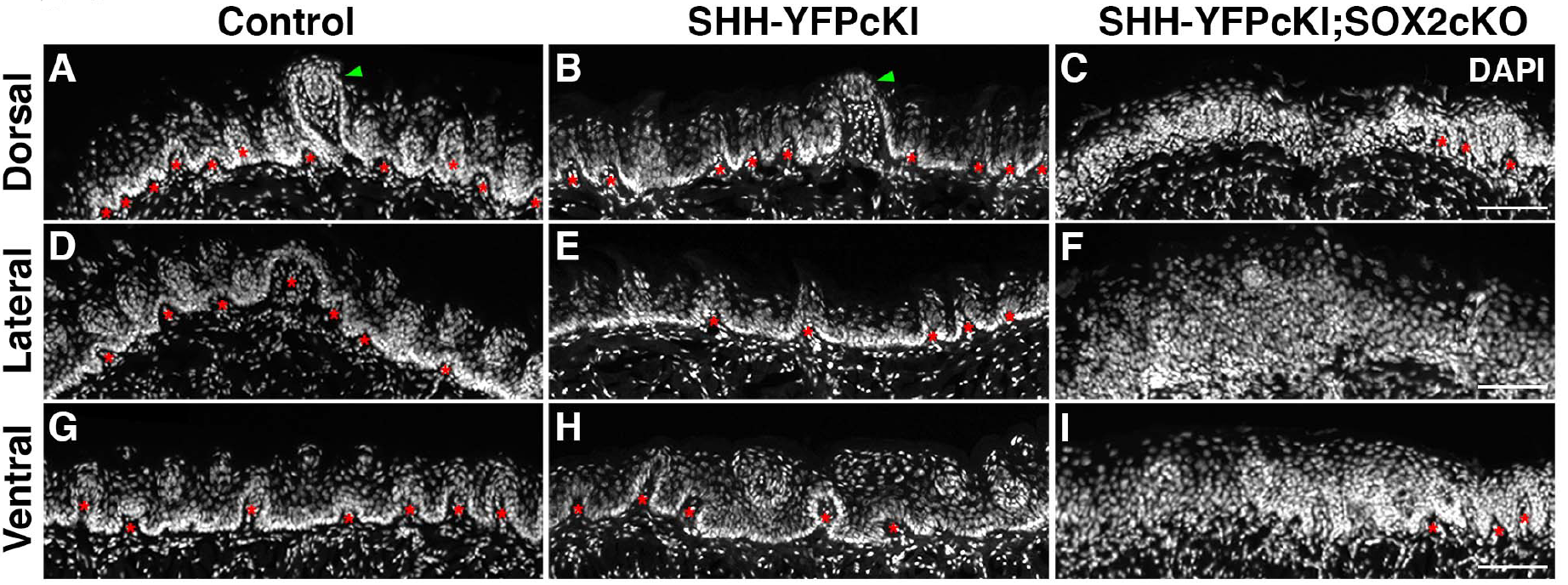
SHH overexpression in SOX2cKO lingual progenitors leads to disorganized epithelia in all regions of the tongue. (A, D, G) Dorsal (A), lateral (D) and ventral (G) epithelium, shown in tranverse sections of control tongue, is composed of non-taste epithelium interspersed with large and small epithelial invaginations (red asterisks) that form part of filiform papillae and fungiform papillae (green arrowhead). Tightly packed basal epithelial cells (Dapi, white) delimit the border between epithelium and lamina propria (lingual mesenchyme). (B, E, H) SHH overexpression in K14^+^ progenitors (SHHcKI) does not affect lingual epithelial morphology; FFP (green arrowhead) and non-taste epithelial invaginations (red asterisks). (C, F, I) All regions of the tongue in SHHcKI;SOX2cKO mice have distorted epithelial architecture, including marked reduction of epithelial invaginations (red asterisks), thickening of the epithelium, and an indistinct border betweent lingual epithelium and the lamina propria. Scale bars = 100 μm. Nuclei are counterstained with DAPI.

